# Dscam1 Controls Presynaptic Terminal Size to Regulate Synaptic Structure and Function in a Central Motor Circuit

**DOI:** 10.1101/2025.09.27.678979

**Authors:** Casey Spencer, Sabrina Jara, Rodney Murphey

## Abstract

The *Drosophila* giant fiber system is a well-characterized escape circuit that enables precise investigation of synaptic development and function. Here, we examined the role of the cell adhesion molecule Dscam1 in shaping presynaptic architecture and supporting circuit performance. Using RNAi-mediated knockdown of Dscam1 in giant fiber interneurons, we observed reduced presynaptic terminal volume, smaller synapse interfaces, and reduced levels of electrical (Shak-B) and chemical (Bruchpilot) synaptic proteins. Despite this reduction in overall abundance, protein localization and density remained unchanged. Functionally, Dscam1 knockdown impaired circuit performance, both increasing latency and reducing success at high frequency demand. Correlation and principal component analysis of our data set show that anatomical variables, particularly presynaptic terminal size, best predicted circuit output. These findings identify Dscam1 as a key developmental regulator of presynaptic terminal size, linking early axon terminal elaboration to adult synaptic function in a central motor circuit. More broadly, our results exemplify a volumetric scaling model where available presynaptic area constrains the distribution of synaptic proteins, both electrical and chemical.

**Significance Statement:** Proper circuit function requires not only the presence of synaptic proteins but also sufficient synaptic space to accommodate them. This study shows that Dscam1 is necessary for normal presynaptic terminal size in a central motor circuit, and when knocked down, constrains synapse size, functional protein content, and subsequently synapse function. These findings highlight a structural mechanism by which developmental regulators can set the capacity for mature synaptic transmission. More generally, our results provide a framework for investigating whether mammalian DSCAM plays a similar role in defining presynaptic capacity and ensuring robust circuit function.

## Introduction

Neural circuit output is shaped not only by patterns of connectivity but also by the architecture of individual synapses. Presynaptic terminal size can influence synaptic efficacy by affecting release probability, transmission capability, and the abundance of active zone proteins (Holderith et al., 2016; Lu et al., 2016). However, proportional scaling between synaptic architecture and function is not always observed; for example, altering protein expression at presynaptic terminals does not necessarily yield proportional changes in circuit output (Goel et al., 2019). The developmental mechanisms that govern presynaptic terminal morphology, as well as the diverse effects of different presynaptic architectures on circuit function, therefore remain incompletely understood (Rollenhagen and Lübke, 2006; Van Vactor and Sigrist, 2017). Understanding how presynaptic terminal size is regulated and how synaptic proteins scale with terminal architecture is thus critical for linking synaptic structure to circuit-level function.

The *Drosophila melanogaster* giant fiber system provides a powerful model for dissecting these relationships. The giant fiber system is a well-characterized escape circuit that rapidly transmits sensory information to flight and jump muscles (Allen et al., 2006). Within the pathway, a mixed electrical and chemical synapse connects the giant fibers (GFs) to the tergotrochanteral motor neurons (TTMns), enabling both rapid signal conduction through gap junctions and chemical transmission (Phelan et al., 1996; Allen and Murphey, 2007). The accessibility of this circuit for both genetic manipulation and direct physiological measurement makes it uniquely suited for investigating how structural features of a presynaptic terminal relate to functional output. Dual-mode analysis, combining confocal quantification of identified synapse architecture and proteins with intracellular recordings from the downstream muscle, provides an opportunity to study central synaptic development with precision and *in vivo* relevance.

Down syndrome cell adhesion molecule 1 (Dscam1) is an immunoglobulin superfamily protein that has been extensively studied for its roles in neural development. Through alternative splicing, Dscam1 can generate tens of thousands of isoforms, allowing individual neurons to express unique combinations that mediate highly specific cell-cell recognition (Schmucker et al., 2000; Sun et al., 2013; Williams et al., 2022). Dscam1 is essential for neurite self-avoidance, ensuring that sister branches of the same neuron do not cross or fasciculate, and it contributes to attractive axon guidance and branch segregation during development (Matthews et al., 2007; Millard et al., 2010). While its role in patterning dendritic arbors and shaping peripheral sensory and motor connections is well established, far less is known about its contributions to synapse formation within the central nervous system and within distinct cell types (Wilhelm et al., 2022). Furthermore, analogous roles for DSCAM family members in vertebrate synapse formation have been observed in the retina and cerebellum, suggesting conserved functions in synaptic assembly and function (Fuerst et al., 2009; Dewa et al., 2024). Nonetheless, the role of Dscam1 in presynaptic terminal development and its role in the function of mixed central synapses remains underexplored.

In particular, it remains unclear whether Dscam1 regulates the formation or maturation of presynaptic terminals in central interneurons. The influence of Dscam1 on the molecular composition of mixed synapses, such as the relative abundance or spatial organization of proteins mediating electrical coupling or chemical release, has not been thoroughly investigated. This gap in knowledge is further compounded by the lack of experimental systems that allow for simultaneous, quantitative assessment of both synaptic structure and corresponding physiological output *in vivo*. Without such systems, it is challenging to determine whether observed morphological changes translate into meaningful differences in circuit performance.

Here, we use the genetically tractable giant fiber system to investigate the role of Dscam1 in the development of a large, identified presynaptic terminal and its functional relationship to circuit physiology. By combining targeted RNAi-mediated knockdown of Dscam1 in GF interneurons with high-resolution confocal imaging and electrophysiological recordings, we quantify how reductions in Dscam1 expression affect terminal size, the localization of synaptic proteins, and the success of signal transmission. This approach provides a direct test of whether a cell adhesion molecule known for mediating axonal patterning also plays a central role in scaling and organizing presynaptic terminals to support reliable high-speed circuit function.

## Methods

### Fly Husbandry and Genetics

Drosophila melanogaster were reared at 25 °C under a 12-hour light/12-hour dark cycle on standard cornmeal-agar media. Adult female flies aged 1-5 days post-eclosion were used for all adult experiments. Pupal growth cone experiments were performed at 24 hours after puparium formation. The UAS-Gal4 system was used to fluorescently label the Giant Fiber (GF) interneurons and drive RNAi expression. The GF-specific Gal4 line, R91H05 (BDSC #40594), was recombined with UAS-mCD8::GFP (provided by Tanja Godenschwege, Florida Atlantic University) to allow for simultaneous labeling and manipulation. Two UAS-RNAi constructs targeting Dscam1 were used: P{KK108835}VIE-260B (VDRC #108835), referred to as Dscam1RNAi (II), and P{GD14362}v36233 (VDRC #36233), referred to as Dscam1RNAi (III). For adult experiments, flies carrying the RNAi constructs were crossed to flies carrying R91H05-Gal4::GFP, and siblings carrying the CyO balancer (but not the RNAi transgene) served as controls. Test genotypes: UAS-Dscam1RNAi (II)/+; R91H05-Gal4::GFP/+ and UAS-Dscam1RNAi (III)/R91H05-Gal4::GFP. Control genotype: CyO/+; R91H05-Gal4::GFP/+.

For pupal experiments, flies were crossed to a Gla, Bc balancer stock to identify animals lacking the RNAi construct. Test genotype: UAS-Dscam1RNAi (II)/+; R91H05-Gal4::GFP/+. Control genotype: Gla,Bc/+; R91H05-Gal4::GFP/+.

### Electrophysiology

Flies were anesthetized with CO₂ and mounted dorsal side up on dental wax. Mounted flies were positioned on a stage under an Amscope stereomicroscope illuminated with white LED lighting. Two glass microelectrodes (∼30 MΩ tip resistance) were pulled, filled with O’Dowd’s dissection saline, and connected via silver wire to a Getting Model 5A amplifier mounted on 3D hydraulic micromanipulators. Electrodes were inserted into the tergotrochanteral (TTM) muscles. Two sharpened tungsten steel stimulating electrodes and one sharpened tungsten steel reference electrode were mounted on 3D micromanipulators and positioned in the brain and abdomen, respectively. Stimulating and reference electrodes were connected to a Grass Instruments isolation unit and an S48 stimulator. Stimuli were delivered at threshold intensity sufficient to elicit GF responses. Amplified (10×) TTM recordings were routed through a Hitachi digital storage oscilloscope and into an Axon Digidata 1440A data acquisition system. Data were recorded and analyzed in ClampX software (Molecular Devices). Following testing, flies were placed under a moistened covered dish to prevent desiccation.

### Tissue Dissections, Immunohistochemistry, Dye-fills, and Imaging

Following electrophysiology, a 10 kDa tetramethylrhodamine dye (Molecular Probes, D1817) solution was backfilled into broken-glass electrodes, mounted in 3D micromanipulators, and inserted into the TTM muscles. Muscles and TTMns were allowed to fill for ∼5 min. Flies were then transferred to a Sylgard-coated dissection dish, and the ventral nerve cord (VNC) was exposed within the thorax (Boerner and Godenschwege, 2011). Tissue was fixed in ice-cold 4% paraformaldehyde (PFA) for 30 min at room temperature, washed in 1× PBS for 30 min, and permeabilized in 0.5% Triton X-100 in PBS for 45 min on an orbital shaker at room temperature. Primary antibodies, prepared in 0.3% Triton X-100 in PBS with 3% BSA, were incubated at 4 °C for 48 h. Samples were washed in PBS for 3-4 h before incubation in secondary antibodies (in PBS) at 4 °C for 24 h, followed by another 3-4 h wash. Tissues were dehydrated through an ethanol series (50%, 70%, 90%, 100%; 8 min each) and cleared in methyl salicylate. Samples were mounted on glass slides with metal spacers, covered with a coverslip, and imaged within 24 h. Samples were imaged on an upright Nikon Eclipse Ti microscope with an A1R detector using a 60× oil-immersion objective (NA 1.4). Laser excitation wavelengths were 488, 561, and 640 nm.

Primary antibodies: mouse anti-Brp (DSHB-NC82, 1:30), mouse anti-GFP (Invitrogen A11120, 1:500), rabbit anti-GFP (Invitrogen A11122, 1:500), rabbit anti-Shak-B (Biomatik, 1:250). The Shak-B antibody was custom-ordered from Biomatik using a previously published sequence targeting all isoforms (CQHHRVPGLKGEIQD) (Phelan et al., 1996; Ammer et al., 2022).

Secondary antibodies: goat anti-mouse 488 (Jackson Immuno 115-545-003, 1:250), goat anti-mouse 647 (Jackson Immuno 115-605-003, 1:250), goat anti-rabbit Cy5 (Jackson Immuno 111-175-144, 1:250).

### Image Analysis and Statistics

Confocal image stacks were converted from *.NIS* format to *.IMS* and opened in Imaris 10.2.0 (Oxford Instruments). The mCD8::GFP (488 nm) fluorescence channel was thresholded and surfaced to best fit the GF axons. The surface was then horizontally segmented to isolate the GF axon terminal on each side of the circuit. The 488 nm (mCD8::GFP) and 561 nm (tetramethylrhodamine) channels were opened in the Imaris colocalization tool, thresholded, and used to generate a new fluorescence channel representing the GF-TTMn synapse interface. A surface of this interface was created. Shak-B or Brp immunolabeling was masked within either the GF terminals or the synapse interface to generate separate fluorescence channels. Each masked channel was thresholded and surfaced to calculate the volumes of: GF axon terminals; GF-TTMn synapse interfaces; Shak-B within GF terminals; Shak-B within synapse interfaces; Brp within GF terminals; Brp within synapse interfaces. Volumes were measured independently for the left and right sides of the circuit in each preparation. Image analysis was kept blinded until the results were obtained. Statistical tests were selected based on data normality and the number of genotypes compared. For three-group comparisons, a Kruskal-Wallis test followed by Dunn’s post hoc test was used. For two-group comparisons, a nonparametric t-test (Mann-Whitney U) was applied. Principal component analysis was performed using scaled anatomical and physiological variables to reduce dimensionality and identify shared variance between datasets. All statistical analyses and graph generation were performed in R (v2024.04.02).

### Growth Cone Visualization

The CNS of developing pupae at 24 hours after pupation onset was dissected in ice-cold O’Dowd’s saline and transferred to poly-lysine-coated glass slides. A plastic ring was mounted onto each slide using Vaseline to contain solutions. Tissues were fixed for 30 min in ice-cold 4% PFA, washed with 1× PBS, and processed for immunohistochemistry as described above, using antibodies to enhance mCD8::GFP fluorescence. Samples were dehydrated, mounted in methyl salicylate under a glass coverslip, and sealed with nail polish.

Samples were imaged on a Nikon Eclipse Ti2 with an Andor Dragonfly 200 spinning disk confocal using a 60× oil immersion objective (NA 1.4). Confocal stacks were analyzed in FIJI: an average intensity projection was thresholded to best fit, and an ROI was drawn around the posterior growth cone region. Fluorescence within the ROI was masked, and growth cone area was quantified for each genotype. Image analysis was kept blinded until the results were obtained.

## Results

### Dscam1 knockdown in the GFs disrupts escape circuit function

The physiology of the *Drosophila* giant fiber system has been extensively characterized. The GFs form mixed electrical and chemical synapses with downstream targets to mediate the rapid escape response (Allen et al., 2006; Uthaman et al., 2008; Allen and Godenschwege, 2010; Boerner and Godenschwege, 2011). In this study, we focused our investigation on the synapse between the GF axon and the medial dendrite of the tergotrochanteral (TTMn) motor neuron within the ventral nerve cord (VNC) (Fig. 1A).

**Figure 1.**
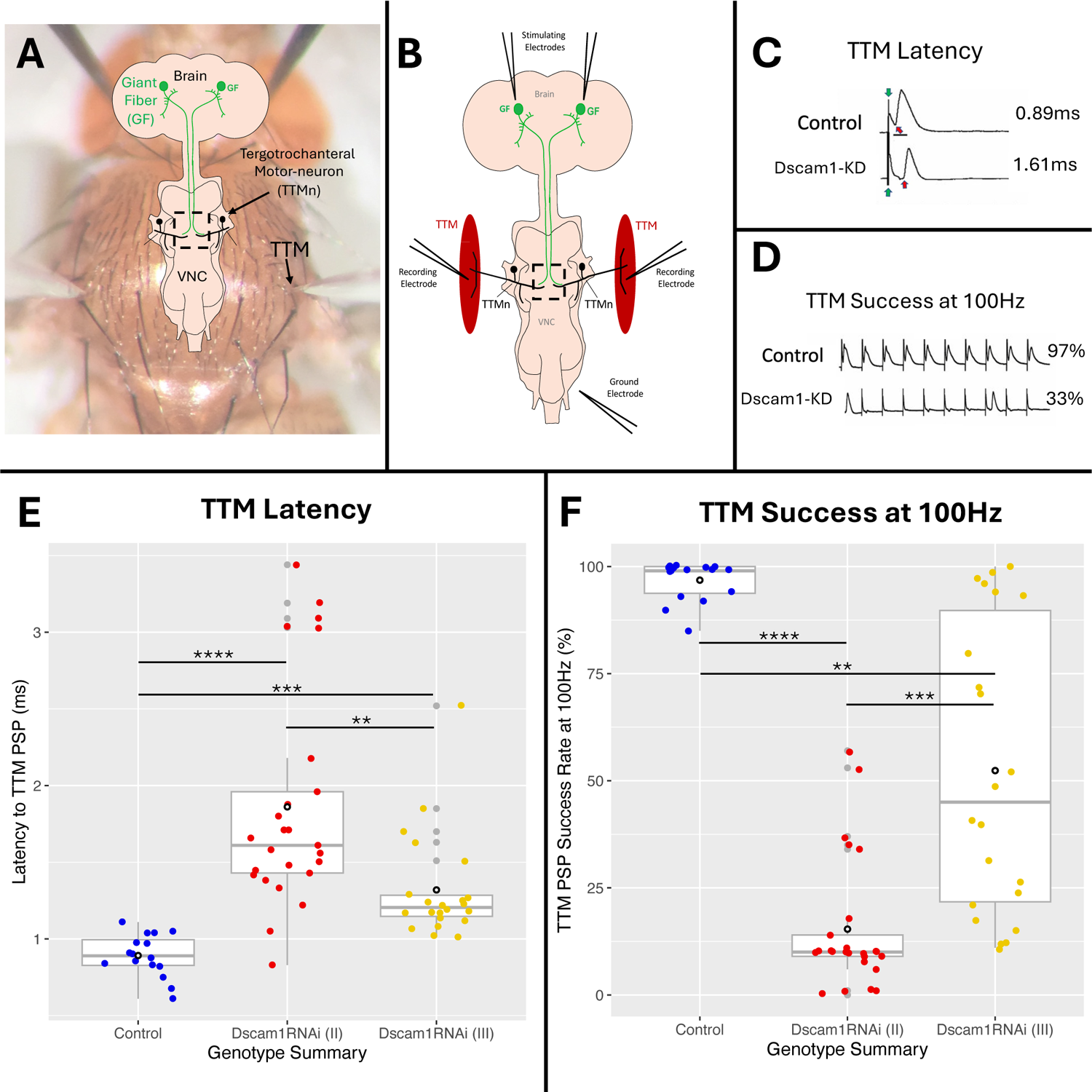
Knockdown of Dscam1 in the GF interneurons impairs circuit latency and success rate. A, Schematic of the Drosophila central nervous system and giant fiber system, shown relative to adult body anatomy. Each giant fiber (GF) axon descends from the brain into the ventral nerve cord (VNC) and forms a mixed electrochemical synapse onto the medial dendrite of the ipsilateral tergotrochanteral motor neuron (TTMn). TTMn axons innervate the tergotrochanteral muscles (TTM), which mediate the jump response during escape behavior. B, Schematic of the electrophysiology setup. Tungsten electrodes are placed in the brain to stimulate the GFs, while glass microelectrodes are inserted into the TTM to record postsynaptic potentials (PSPs). The stimulation circuit is grounded via a tungsten electrode in the abdomen; recording amplifiers use internal AC reference. C, Representative PSP recordings showing increased latency following GF-specific Dscam1 knockdown. Control flies exhibit a latency of 0.89 ms, while flies with Dscam1 knockdown show a mean latency of 1.61 ms (combined data from both RNAi lines). D, Representative traces showing reduced PSP success at 100 Hz following Dscam1 knockdown. Control flies maintain robust PSPs across stimuli (∼97% success), while knockdown animals show reduced circuit success (∼33% success). E, Box-and-whisker plot quantifying TTM PSP latency. Dscam1 knockdown in GFs significantly prolongs latency compared to controls (mean = 0.89 ms; *n* = 16), with Dscam1RNAi (II) showing a stronger effect (mean = 1.86 ms; *n* = 25; *p* = 2.34 × 10⁻⁹) than Dscam1RNAi (III) (mean = 1.32 ms; *n* = 22; *p* = 3.46 × 10⁻⁴). The two RNAi lines also differ from one another (*p* = 9.44 × 10⁻³). F, Box-and-whisker plot quantifying TTM PSP success rate at 100 Hz. Dscam1 knockdown in GFs significantly impairs circuit success relative to controls (mean = 96.81%; *n* = 16), with Dscam1RNAi (II) exhibiting a more severe reduction (mean = 15.36%; *n* = 25; *p* = 3.83 × 10⁻¹⁰) than Dscam1RNAi (III) (mean = 52.36%; *n* = 22; *p* = 3.32 × 10⁻³). The RNAi lines also differ significantly (*p* = 2.75 × 10⁻⁴). Each dot represents a single circuit side and resulting TTM response; black circle indicates the group mean. Gray dots denote statistical outliers. ns = not significant; * < 0.05; ** < 0.01; *** < 0.001; **** < 0.0001. Statistical analysis was performed using a Kruskal-Wallis test with Dunn’s post hoc test.

Using previously reported methods (Allen and Godenschwege, 2010), stimulating electrodes were placed into the brain of the adult female fly to excite the GFs. Recording electrodes were placed in each TTM, and the recordings yield a post-synaptic potential (PSP) (Fig. 1B). We quantified two physiological parameters to assess circuit function: (1) latency, defined by the time between GF stimulation and TTM response, and (2) PSP success at 100 Hz, the success rate, of the circuit firing under high-frequency stimulation. In control flies, the TTM displayed a latency of 0.89 ms and a high-probability PSP success of 97% at 100 Hz (Fig. 1C-D).

To determine whether Dscam1 is required for normal circuit function, we knocked down Dscam1 specifically in the GFs using two separate UAS-Dscam1-RNAi constructs (Wilhelm et al., 2022) driven by a strong GF-specific R91H05-Gal4 line (Borgen et al., 2017). Expression of this GF Gal4 driver and subsequent knockdown of Dscam1 in the GFs occurs early in development before the GF axon reaches the TTMn medial dendrite (Lee and Godenschwege, 2015). We assessed the phenotypes from each RNAi line to confirm that the observed effects resulted specifically from Dscam1 knockdown and not from off-target effects. VDRC #108835 is referred to here as Dscam1RNAi (II), and VDRC #36233 as Dscam1RNAi (III). These lines differ in chromosomal insertion site (chromosomes 2 and 3, respectively) and non-overlapping RNAi target region (data not shown). To measure the effects of Dscam1-KD in the GFs, both RNAi lines were expressed under identical conditions and analyzed between 1-5 days post-eclosion. All control animals were genetic siblings of experimental animals, lacking the RNAi construct, and exhibited previously reported wild-type responses (Allen and Godenschwege, 2010).

Knockdown of Dscam1 in the GFs significantly impaired both latency and response success of signal transmission through the circuit (Fig. 1E-F). Dscam1RNAi (II) caused a severe delay in TTM latency (mean = 1.86 ms, *p* = 2.34 × 10⁻⁹), while Dscam1RNAi (III) caused a milder but still significant increase (mean = 1.32 ms, *p* = 3.46 × 10⁻⁴), relative to controls (mean = 0.89 ms). The latency values also differed significantly between the two RNAi lines (*p* = 9.44 × 10⁻³). Similarly, PSP success was reduced following knockdown of Dscam1. Dscam1RNAi (II) animals exhibited severely diminished success rates at 100 Hz stimulation (mean = 15.36%, *p* = 3.83 × 10⁻¹⁰), while Dscam1RNAi (III) animals showed a moderate but significant reduction (mean = 52.36%, *p* = 3.32 × 10⁻³), compared to controls (mean = 96.81%). The difference between RNAi lines was also significant (*p* = 2.75 × 10⁻⁴).

These findings indicate that Dscam1 expression in the GF interneurons is essential for establishing a functional, high-fidelity synapse with the TTMn, and that the severity of circuit disruption varies with RNAi construct, suggesting a dose-dependent or target-region-specific effect. We next examined the structural features of the GF axon terminal in response to Dscam1 knockdown.

### Dscam1 controls synaptic terminal volume of the GF axon

Dscam1 is a known regulator of axon guidance and neurite self-avoidance across multiple model systems (Hattori et al., 2008; Fuerst et al., 2009; Huang et al., 2011; Matthews and Grueber, 2011; Cvetkovska et al., 2013; Kim et al., 2013; Montesinos, 2014; Dascenco et al., 2015; Sachse et al., 2019). The *Drosophila* giant fiber system provides a powerful *in vivo* model to dissect the developmental roles of Dscam1 in synaptic morphology and circuit formation. In wild-type flies, the GF axon projects posteriorly from the brain and bends laterally within the VNC to form a mixed electrical and chemical synapse with the ipsilateral TTMn. This terminal bend defines the GF synaptic terminal and is highly sensitive to Dscam1 knockdown (Fig. 2A-B). Surprisingly, targeted knockdown of Dscam1 in the GFs did not impair gross axon pathfinding: GF axons consistently reached the second thoracic neuromere of the ventral nerve cord (VNC) and formed a synapse (data not shown). However, many GFs failed to elaborate a normal synaptic terminal, and those that did exhibited markedly reduced terminal volume.

**Figure 2.**
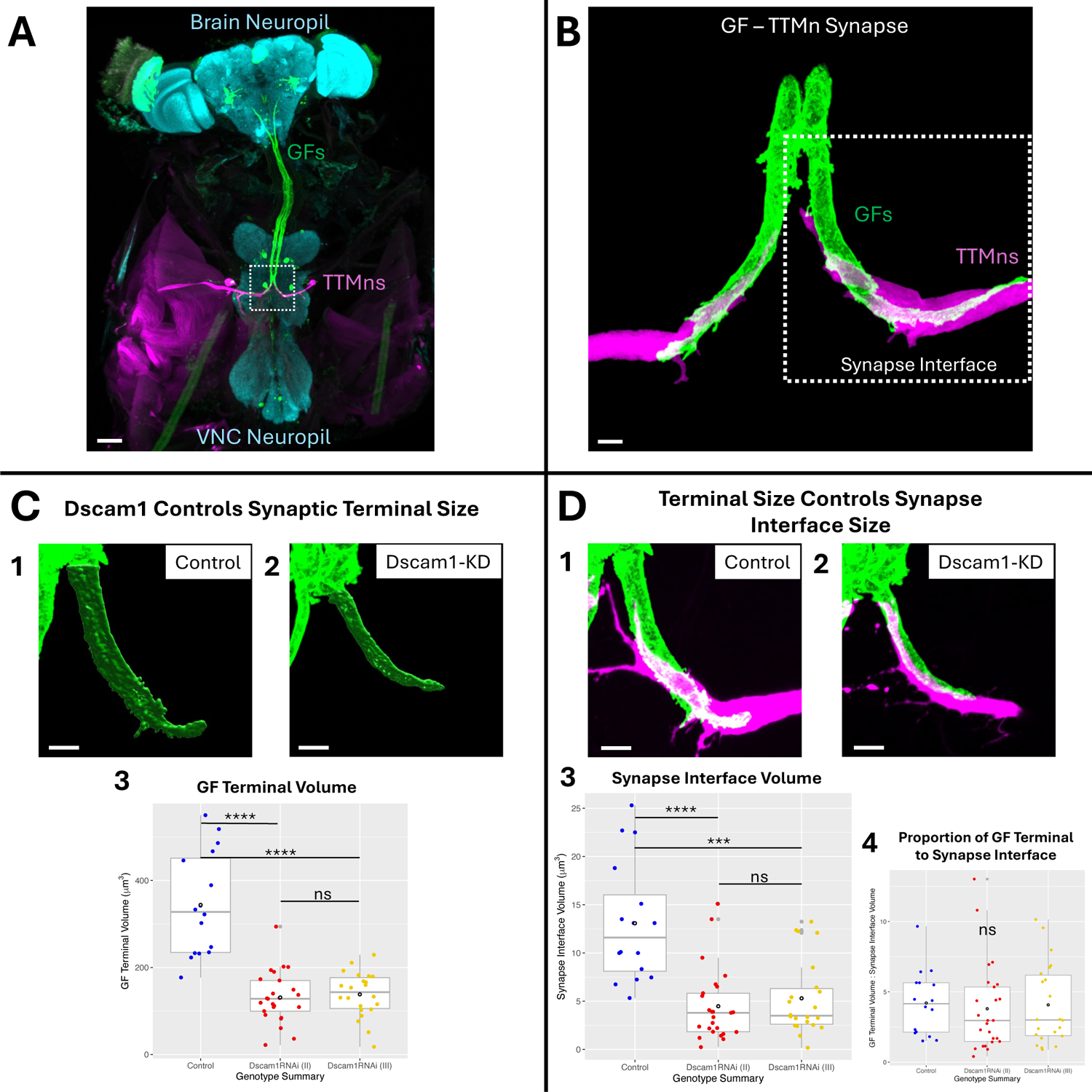
Dscam1 controls GF axon terminal volume and influences synapse size. A, 3D confocal image of the Drosophila CNS (20× objective), with a white dotted box indicating the GF-TTMn synapse. Genetically expressed mCD8::GFP labels GFs (green), rhodamine dye injection labels TTMns (magenta), and Bruchpilot (Brp) antibody labels neuropil (blue). Scale bar = 50 μm. B, 3D confocal image of a control GF-TTMn synapse (60× objective). GFs (green), TTMns (magenta), and their colocalized overlap (white) highlight the synapse interface. The white dotted box outlines the region of interest (ROI) used for quantification. Scale bar = 5 μm. C1-C2, Representative images of GF terminals from control (C1) and Dscam1-KD (C2) animals, showing visibly shorter and thinner terminals. Scale bars = 5 μm. C3, Box-and-whisker plot quantifying GF terminal volume. Dscam1-KD significantly reduces terminal volume compared to controls (mean = 343.88 μm³; *n* = 16). Dscam1RNAi (II) (mean = 131.12 μm³; *n* = 25; *p* = 2.95 × 10⁻⁷) results in a slightly stronger phenotype than Dscam1RNAi (III) (mean = 138.2 μm³; *n* = 22; *p* = 2.8 × 10⁻⁶). Each dot represents one GF terminal; black circle: group mean; grey dots: statistical outliers. Kruskal-Wallis test with Dunn’s post hoc test. D1-D2, Representative images of GF-TTMn synapses from control (D1) and Dscam1-KD (D2) animals showing visibly reduced synapse interface size in KD animals. Scale bars = 5 μm. D3, Box-and-whisker plot quantifying synapse interface volume. Dscam1-KD significantly reduces synapse interface volume relative to controls (mean = 13.08 μm³; *n* = 16), with Dscam1RNAi (II) (mean = 4.48 μm³; *n* = 25; *p* = 1.49 × 10⁻⁵) showing a slightly stronger effect than Dscam1RNAi (III) (mean = 5.30 μm³; *n* = 22; *p* = 2.04 × 10⁻⁴). Each dot represents a single synapse interface; black circle: group mean; grey dots: statistical outliers. Kruskal-Wallis test with Dunn’s post hoc test. D4, Proportional analysis of the GF synaptic terminal volume occupied by the synapse interface reveals no significant differences across genotypes (∼4%; *p* > 0.05), suggesting that interface size scales proportionally with terminal volume and is independent of Dscam1 abundance.

Dscam1 knockdown (Dscam1-KD) resulted in a variability of presynaptic morphological defects, including shortened and thinned synaptic terminals. To label the GF membrane and drive RNAi expression, we used a recombined R91H05-Gal4 line expressing membrane-bound GFP (UAS-mCD8::GFP). Confocal Z-stacks of GFP-labeled GF axons were acquired using point-scan microscopy and processed in Imaris software for volumetric analysis. Each GF axon terminal fluorescence was thresholded to best-fit, and surfaced to yield a volume. The surface of each sample was cut at the same anatomical landmark as marked by the lateral axon bend (Fig. 2C1-C2). Quantification revealed that GF synaptic terminal volume was significantly reduced in Dscam1-KD animals compared to controls (mean = 343.88 μm³). Dscam1RNAi (II) resulted in the strongest reduction (mean = 131.12 μm³; *p* = 2.95 × 10⁻⁷), with Dscam1RNAi (III) exhibiting a slightly milder effect (mean = 138.2 μm³; *p* = 2.8 × 10⁻⁶) (Fig. 2C1-C3). These findings establish Dscam1 as a key regulator of presynaptic terminal development in a central interneuron. While previous Dscam1 loss-of-function studies have reported reduced axonal branching and arbor size in sensory neurons (Kim et al., 2013; Chen et al., 2022), our findings provide the first quantitative evidence that Dscam1 controls specifically the synaptic terminal volume in an interneuron. We next sought to identify the subcellular changes to the synapse made between the GF and the TTMn.

### Synaptic terminal volume predicts GF-TTMn synapse interface size

To evaluate how Dscam1 knockdown affects synaptic connectivity between the GF and TTMn, we quantified the region of membrane apposition between these two cells, referred to here as the GF-TTMn synapse interface, using confocal microscopy. To effectively visualize the synapse interface, we simultaneously labeled the GF axon and the TTMn dendrite. The GF axon was genetically labeled with membrane-bound GFP, while the TTMn was backfilled by injecting tetramethylrhodamine dye (10 kDa) into the TTM muscle, which retrogradely labeled the TTMn medial dendrite (Fig. 2B). The synapse interface was defined as the region of colocalization between the pre- and post-synaptic fluorescent signals, captured using point-scan confocal microscopy and reconstructed in Imaris software. Although EM reveals synaptic contacts as 2D zones, confocal imaging enables 3D volumetric quantification following colocalization thresholding and surfacing. The validity of this approach is supported by prior studies showing that the synapse can be quantitatively analyzed in confocal Z-stacks, paired with the labeling of synaptic structures such as Synapsin-1 and confirmed via correlative EM (Hohensee et al., 2008).

Compared to controls (mean = 13.08 μm³), Dscam1 knockdown significantly reduced synapse interface volume. Dscam1RNAi (II) reduced interface volume to 4.48 μm³ (*p* = 1.49 × 10⁻⁵), and Dscam1RNAi (III) to 5.30 μm³ (*p* = 2.04 × 10⁻⁴) (Fig. 2D3). Despite this reduction in absolute size, the proportion of each GF terminal devoted to the synapse interface (∼4%) remained unchanged across genotypes (*p* > 0.05; Fig. 2D4).

These results suggest that Dscam1 regulates terminal growth and that synapse interface scales directly with presynaptic terminal volume. This interpretation aligns with our physiological data: smaller terminals form smaller synapses and exhibit impaired synaptic output. To further investigate how Dscam1 influences synaptic function, we next examined the localization and abundance of key synaptic proteins, Shak-B (electrical) and Brp (chemical).

### Dscam1 is necessary for gap junction protein abundance

To determine whether Dscam1 affects the abundance or distribution of synaptic components, we examined the localization of the gap junction protein Shaking-B (Shak-B), encoded by the shakB gene (Phelan et al., 1996), within both the GF synaptic terminal and the GF-TTMn synapse interface. We used a custom primary antibody directed against all isoforms of Shak-B, followed by fluorescently tagged secondary antibodies to visualize Shak-B in confocal z-stacks. Using Imaris software, we quantified the colocalization of Shak-B immunofluorescence with GFP-labeled GF terminals, thresholded/surfaced their fluorescence, and measured the Shak-B volume (Fig. 3A).

**Figure 3.**
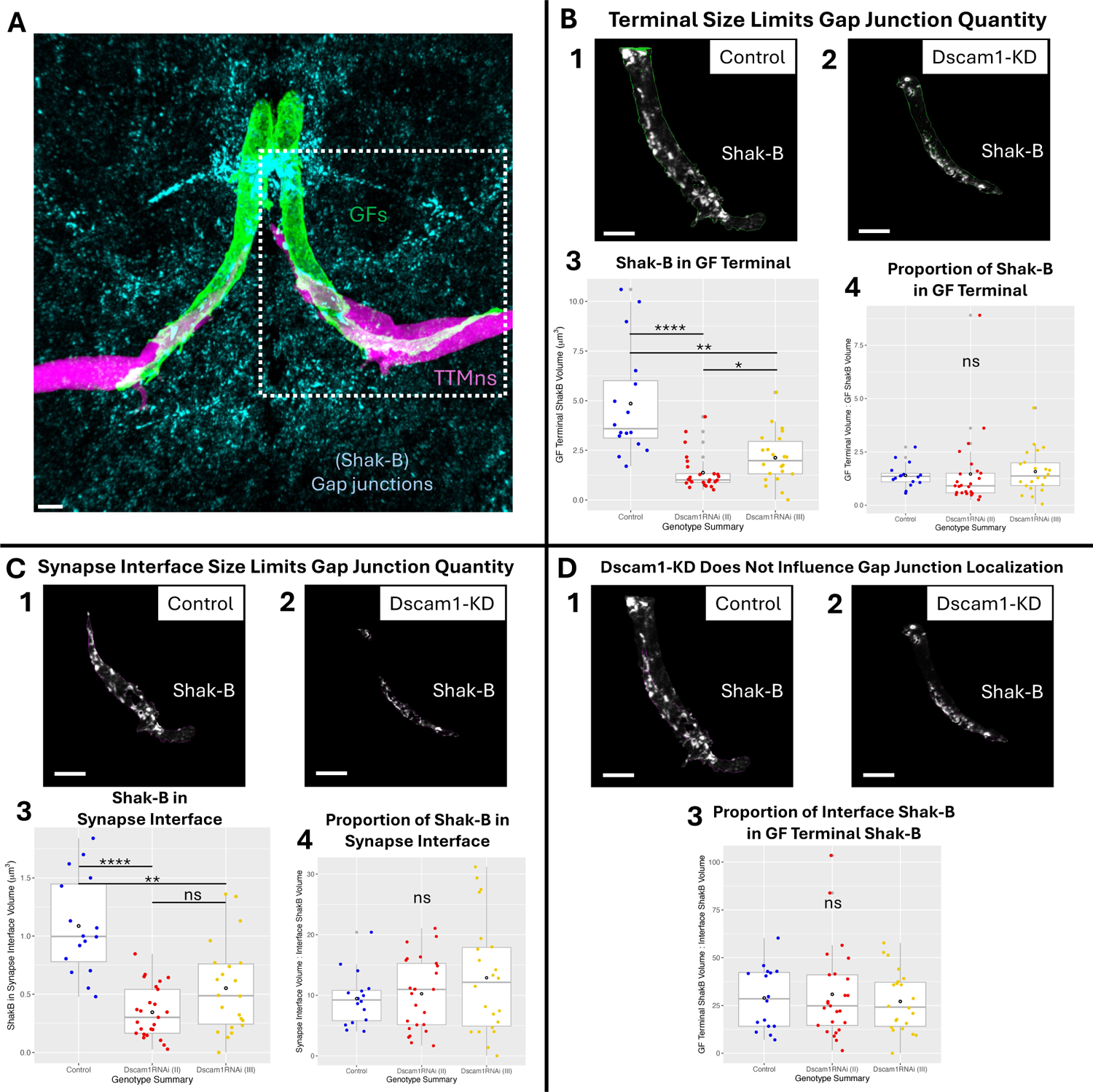
Dscam1-KD in the GFs reduces the quantity of gap junctions at the GF-TTMn synapse. A, 3D confocal image of a control GF-TTMn synapse interface showing Shak-B gap junctions (60× objective). GFs (green) are labeled with genetically expressed mCD8::GFP, TTMns (magenta) with rhodamine dye injection, and gap junctions (light blue) with anti-Shak-B immunolabeling. The white dotted box outlines the region of interest (ROI) analyzed in subsequent panels. Scale bar = 5 μm. B1-B2, Representative 3D confocal images of Shak-B localization within control (B1) and Dscam1-KD (B2) GF terminals, showing reduced Shak-B volume following KD. Scale bars = 5 μm. B3, Box-and-whisker plot quantifying Shak-B volume within GF terminals. Dscam1-KD significantly reduces Shak-B levels compared to controls (mean = 4.85 μm³; *n* = 16). Dscam1RNAi (II) (mean = 1.37 μm³; *n* = 25; *p* = 5.79 × 10⁻⁷) results in a slightly stronger phenotype than Dscam1RNAi (III) (mean = 2.13 μm³; *n* = 22; *p* = 3.47 × 10⁻³). The two KD constructs also differ from each other (*p* = 2.28 × 10⁻²). Each dot represents a single terminal; black circle: group mean; grey dots: statistical outliers. Kruskal-Wallis test with Dunn’s post hoc test. B4, The proportion of Shak-B volume within the GF axon terminal remains unchanged across genotypes (1.39–1.57%; *p* > 0.05), suggesting that gap junction scaling is proportional to terminal size and not directly regulated by Dscam1 abundance. C1-C2, Representative images of Shak-B fluorescence distribution at the GF-TTMn synapse interface in control (C1) and Dscam1-KD (C2) animals. Dscam1-KD shows reduced Shak-B puncta. Scale bars = 5 μm. C3, Box-and-whisker plot quantifying Shak-B volume within the synapse interface. Dscam1-KD significantly reduces Shak-B compared to controls (mean = 1.09 μm³; *n* = 16), with Dscam1RNAi (II) (mean = 0.35 μm³; *n* = 25; *p* = 1.21 × 10⁻⁶) exhibiting a slightly stronger phenotype than Dscam1RNAi (III) (mean = 0.55 μm³; *n* = 22; *p* = 1.36 × 10⁻³). No significant differences were found between the KD constructs (*p* > 0.05). Each dot represents a single synapse; black circle: group mean; grey dots: statistical outliers. Kruskal-Wallis test with Dunn’s post hoc test. C4, The proportion of Shak-B volume within the synapse interface is unchanged across genotypes (9.5–12.9%; *p* > 0.05), further supporting proportional gap junction scaling relative to terminal size, independent of Dscam1. D1-D2, Representative images showing GF–TTMn synapse interface surfaces and Shak-B distribution within GF terminals from control (D1) and Dscam1-KD (D2) animals. Scale bars = 5 μm. D3, Box-and-whisker plot showing the proportion of total GF Shak-B localized to the synapse interface. No significant differences are observed across genotypes (27.1–30.9%; *p* > 0.05), indicating that Dscam1 does not alter gap junction localization but instead regulates overall quantity via terminal size.

Dscam1 knockdown (Dscam1-KD) significantly reduced presynaptic terminal size (as shown previously in Fig. 2). In parallel, Shak-B absolute abundance within the GF synaptic terminal was also significantly reduced: control animals exhibited a mean Shak-B volume of 4.85 μm³, while Dscam1RNAi (II) and Dscam1RNAi (III) knockdown flies showed reduced levels of 1.37 μm³ and 2.13 μm³, respectively (Fig. 3B1-B3; *p* = 5.79 × 10⁻⁷ for RNAi II vs. control; *p* = 3.47 × 10⁻³ for RNAi III vs. control). The two RNAi constructs also differed significantly from one another (*p* = 2.28 × 10⁻²). However, the proportion of the GF terminal volume occupied by Shak-B remained unchanged across genotypes (control = 1.39%, RNAi II = 1.57%, RNAi III = 1.53%; *p* > 0.05) (Fig. 3B4). These findings suggest that Dscam1 does not directly regulate Shak-B protein levels but instead constrains its abundance indirectly by limiting synaptic terminal size.

To evaluate gap junction abundance at the site of the synapse, we next examined Shak-B volume specifically within the GF-TTMn synapse interface surface volume. Dscam1-KD significantly reduced Shak-B volume at the synapse interface compared to controls (mean = 1.09 μm³ in control, 0.35 μm³ in RNAi II, 0.55 μm³ in RNAi III; *p* = 1.21 × 10⁻⁶ for RNAi II vs. control; *p* = 1.36 × 10⁻³ for RNAi III vs. control) (Fig. 3C1-C3). No significant difference was observed between the two RNAi lines (*p* > 0.05). The proportion of the synapse interface volume occupied by Shak-B remained consistent across groups (control = 12.9%, RNAi II = 9.5%, RNAi III = 10.3%; *p* > 0.05) (Fig. 3C4), further supporting the conclusion that Shak-B quantity scales with synapse size rather than being directly controlled by Dscam1.

Finally, we assessed whether Dscam1 regulates the targeting or localization of Shak-B by comparing the proportion of total Shak-B within the GF terminal that localizes to the synapse interface. Across genotypes, ∼27-31% of terminal Shak-B was localized to the synapse interface (control = 30.9%, RNAi II = 27.1%, RNAi III = 29.2%; *p* > 0.05) (Fig. 3D1-D3), indicating that Dscam1 knockdown does not disrupt the spatial targeting of gap junctions. Together, these findings support a model in which Dscam1 indirectly regulates synaptic protein abundance by limiting the available terminal volume, while localization mechanisms remain intact.

### Synaptic architecture predicts circuit function

To evaluate how Dscam1-dependent changes in synaptic structure relate to physiological output, we directly correlated presynaptic anatomical features with circuit performance using a dual-mode data set in which both structure and function were assessed in the same animal. Pearson correlation analyses revealed that larger GF synaptic terminals were positively associated with increased PSP success rates at 100 Hz (R = 0.64, *p* = 1.8 × 10⁻⁸; Fig. 4A1). Likewise, larger GF-TTMn synapse interfaces correlated with improved PSP success (R = 0.47, *p* = 9.4 × 10⁻⁵; Fig. 4A2). We observed similar relationships for Shak-B gap junction content: both the absolute volume of Shak-B within the GF terminal (R = 0.49, *p* = 4.1 × 10⁻⁵; Fig. 4A3) and at the synapse interface (R = 0.47, *p* = 1.1 × 10⁻⁴; Fig. 4A4) were predictive of higher PSP success. These findings suggest that both terminal size and total gap junction abundance are important predictors of reliable synaptic transmission.

**Figure 4.**
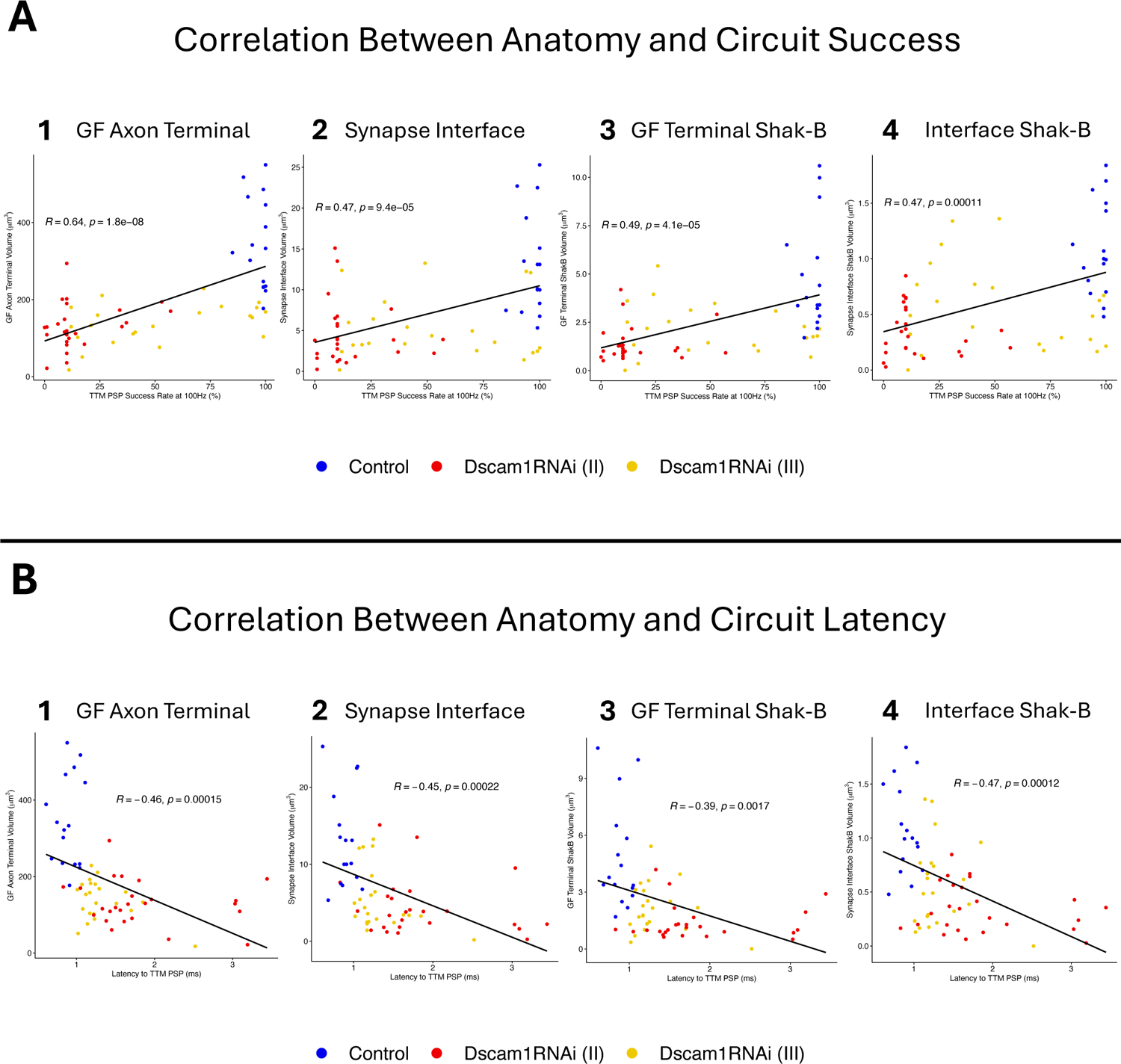
GF-TTMn synaptic structure and gap junction abundance predict circuit performance. A1, Pearson correlation between GF synaptic terminal volume and circuit success at 100Hz. Larger presynaptic terminals are positively associated with higher PSP success rates at 100 Hz (R = 0.64, *p* = 1.8 × 10⁻⁸). Each point represents a single side of the synapse, plotted using paired GF terminal volume and TTM PSP success. A2, Correlation between GF-TTMn synapse interface volume and circuit success. Larger interfaces correlate with improved PSP success (R = 0.47, *p* = 9.4 × 10⁻⁵). A3, Correlation between GF terminal Shak-B volume and circuit success. Greater presynaptic Shak-B content is associated with enhanced PSP success (R = 0.49, *p* = 4.1 × 10⁻⁵). A4, Correlation between synapse interface Shak-B volume and circuit success. Increased Shak-B at the interface corresponds to greater PSP success (R = 0.47, *p* = 1.1 × 10⁻⁴). B1, Pearson correlation between GF synaptic terminal volume and circuit latency. Larger terminals are associated with shorter PSP latencies (R = –0.46, *p* = 1.5 × 10⁻⁴). B2, Correlation between GF-TTMn synapse interface volume and circuit latency. Larger interfaces correlate with faster responses (R = –0.45, *p* = 2.2 × 10⁻⁴). B3, Correlation between GF terminal Shak-B volume and circuit latency. Higher gap junction content in the presynaptic terminal predicts faster response (R = –0.39, *p* = 1.7 × 10⁻³). B4, Correlation between synapse interface Shak-B volume and circuit latency. Greater Shak-B at the synapse is associated with shorter PSP latency (R = –0.47, *p* = 1.2 × 10⁻⁴). Each plot shows one side of the synapse from paired sample data. Positive correlations in panel A reflect improved circuit success with increased anatomical features; negative correlations in panel B reflect improved response speed with increased anatomical features.

We next asked whether these anatomical variables also predicted circuit latency. Correlation analyses demonstrated that larger presynaptic terminals were associated with significantly shorter TTM PSP latencies (R = –0.46, *p* = 1.5 × 10⁻⁴; Fig. 4B1), as were larger synapse interfaces (R = –0.45, *p* = 2.2 × 10⁻⁴; Fig. 4B2). Similarly, greater Shak-B content in the GF terminal (R = –0.39, *p* = 1.7 × 10⁻³; Fig. 4B3) and synapse interface (R = –0.47, *p* = 1.2 × 10⁻⁴; Fig. 4B4) correlated with faster response times, indicating that gap junction abundance influences not only the reliability but also the speed of synaptic transmission.

While these pairwise correlations are informative, they do not account for the interdependence of structural variables. To determine whether combined anatomical features collectively predict synaptic performance, we performed Principal Component Analysis (PCA) on all four structural variables: GF terminal volume, synapse interface volume, GF terminal Shak-B content, and interface-localized Shak-B. Principal Component 1 (PC1) explained 66.1% of the total variance in the dataset and significantly correlated with circuit success (R = 0.638, *p* = 1.86 × 10⁻⁸). Multiple linear regression further revealed that PC1 significantly predicted circuit performance (β = 15.38, *p* = 2.07 × 10⁻⁸), where β reflects the expected increase in circuit success per unit increase in PC1 score, while PC2 did not contribute meaningfully to the model (*p* = 0.38). Together, these findings demonstrate that structural features of the GF-TTMn synapse vary along a common axis and collectively influence circuit output.

These analyses reinforce the model that Dscam1 ensures the development of an appropriately sized presynaptic terminal capable of supporting sufficient interface area and gap junction protein incorporation. Reductions in any of these anatomical variables result in diminished functional performance, linking structural integrity to synaptic output.

### Dscam1 is necessary for chemical synapse abundance

In addition to gap junctions, the GF-TTMn synapse includes chemical synaptic components, including presynaptic active zones that utilize acetylcholine as a neurotransmitter. Although the importance of the chemical component of the GF-TTMn synapse has not been fully elucidated physiologically (Allen et al., 1999; Allen and Murphey, 2007), we investigated possible anatomical changes to the chemical component in response to Dscam1-KD. To determine whether Dscam1 may separately regulate chemical synapse assembly, we quantified the distribution of Bruchpilot (Brp), a core active zone protein, using Brp-specific immunolabeling and volumetric analysis in the GF synaptic terminals and synapse interface (Fig. 5A).

**Figure 5.**
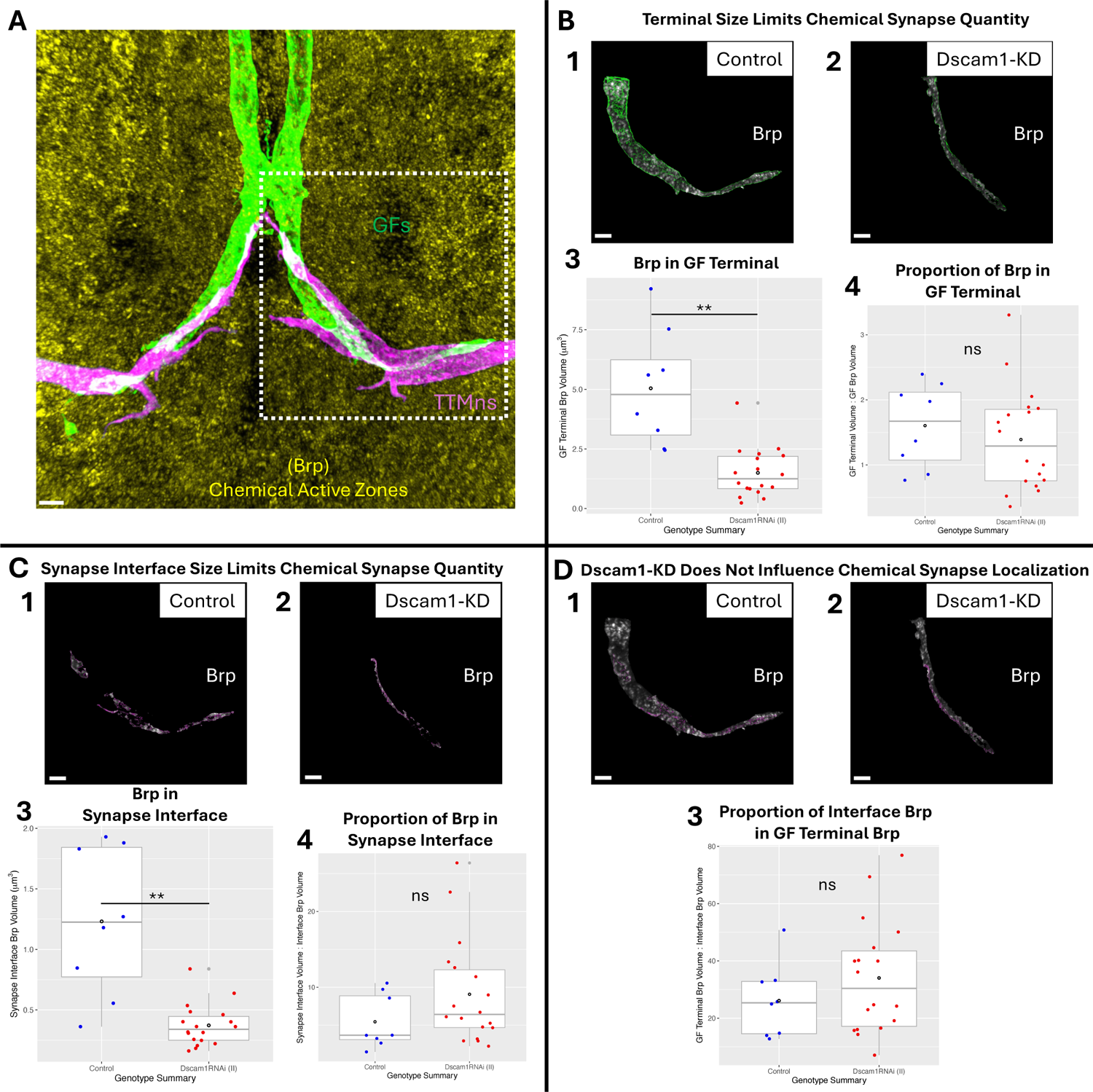
Dscam1-KD in the GFs reduces the quantity of chemical synapses at the GF-TTMn synapse. A, 3D confocal image of the control GF-TTMn synapse interface showing Brp-positive chemical synapses (60× objective). Genetically expressed mCD8::GFP labels the GFs (green), rhodamine dye labels the TTMns (magenta), and Brp-positive active zones are immunolabeled in yellow. The white dotted box indicates the region of interest (ROI) analyzed below. Scale bar = 5 μm. B1-B2, Representative 3D confocal images of Brp fluorescence in control (B1) and Dscam1-KD (B2) GF synaptic terminals. The GF axon terminal surface is outlined in green, and Brp signal is shown in white. Dscam1-KD terminals show reduced Brp content. Scale bar = 4 μm. B3, Box-and-whisker plot quantifying Brp volume within GF terminals. Dscam1RNAi (II) knockdown results in significantly reduced Brp volume compared to controls (Control: mean = 5.05 μm³; *n* = 8; KD: mean = 1.50 μm³; *n* = 18; *p* = 4.15 × 10⁻³; Welch two-sample t-test). B4, The proportion of Brp volume relative to terminal volume is not altered by Dscam1 knockdown (Control: 1.6%; KD: 1.38%; *p* > 0.05), suggesting chemical synapse abundance scales proportionally with terminal size independent of Dscam1 levels. C1-C2, Representative images of Brp fluorescence in control (C1) and Dscam1-KD (C2) GF-TTMn synapse interfaces. Synapse interface volumes are outlined in magenta. Brp signal is reduced in KD animals. Scale bar = 4 μm. C3, Quantification of Brp volume in the synapse interface shows a significant reduction in Dscam1RNAi (II) animals (Control: mean = 1.23 μm³; *n* = 8; KD: mean = 0.37 μm³; *n* = 18; *p* = 5.2 × 10⁻³; Welch two-sample t-test). C4, The proportion of Brp within the interface volume does not differ between genotypes (Control: 5.45%; KD: 9.08%; *p* > 0.05), supporting proportional scaling with structure size. D1-D2, Representative 3D images of GF terminals showing Brp localization within the synapse interface (outlined in magenta) for control (D1) and KD (D2) animals. Scale bar = 4 μm. D3, Quantification of the proportion of GF-terminal Brp localized to the synapse interface shows no difference between genotypes (Control: 26.1%; KD: 34.1%; *p* > 0.05), indicating that Dscam1 does not influence the spatial localization of chemical synapses.

Dscam1 knockdown in the GF significantly reduced Brp volume within the presynaptic terminal (Fig. 5B). Brp content in Dscam1RNAi (II) animals was significantly decreased (mean = 1.50 μm³) relative to controls (mean = 5.05 μm³; *p* = 4.15 × 10⁻³; Fig. 5B3). Despite this reduction in absolute Brp abundance, the proportion of terminal volume occupied by Brp did not differ significantly between genotypes (Control: 1.6%; KD: 1.38%; *p* > 0.05; Fig. 5B4), suggesting that Brp density is preserved and that Dscam1 influences Brp levels via modulation of terminal size.

A similar pattern was observed in the synapse interface (Fig. 5C). Dscam1RNAi (II) animals displayed significantly reduced Brp content at the GF-TTMn synapse interface (mean = 0.37 μm³) compared to controls (mean = 1.23 μm³; *p* = 5.2 × 10⁻³; Fig. 5C3). However, the relative proportion of interface volume occupied by Brp was unchanged across genotypes (Control: 5.45%; KD: 9.08%; *p* > 0.05; Fig. 5C4), further indicating that Brp levels scale with structural dimensions rather than being directly controlled by Dscam1.

Finally, we examined the localization of Brp relative to the synapse interface and found no evidence that Dscam1 alters Brp spatial distribution (Fig. 5D). The proportion of Brp within the GF terminal that localized to the synapse interface was comparable between genotypes (Control: 26.1%; KD: 34.1%; *p* > 0.05; Fig. 5D3). These results mirror those obtained with Shak-B and confirm that while Dscam1 regulates the overall abundance of synaptic components, it does not influence their density or spatial targeting.

Together, these data support a model in which Dscam1 ensures the formation of a presynaptic terminal of sufficient size to incorporate both electrical and chemical synaptic components. While protein density and localization remain stable, overall synapse composition is constrained by synaptic growth, underscoring the structural role of Dscam1 in synapse assembly.

### Dscam1 is required during synaptogenesis to regulate GF growth cone size

Dscam1 has well-established roles in neurite self-avoidance, axon guidance, and growth cone dynamics across diverse neuronal types (Hughes et al., 2007; Soba et al., 2007; Andrews et al., 2008a, 2008b; Hattori et al., 2009; Cvetkovska et al., 2013; Kim et al., 2013; Alavi et al., 2016). To determine whether Dscam1 influences the early structural events that precede synapse formation in the GFs, we examined GF axon morphology at 24 hours after pupation begins. This represents a key developmental stage when axons are actively navigating and initiating contact within the second thoracic neuromere of the ventral nerve cord (Lee and Godenschwege, 2015).

We labeled GF axons with membrane-bound GFP driven by R91H05-Gal4 and imaged the distal axon tips under confocal microscopy. At this stage, filopodial processes were visible, but mature synaptic connections with the tergotrochanteral motor neuron (TTMn) had not yet formed (Fig. 6A-B).

**Figure 6.**
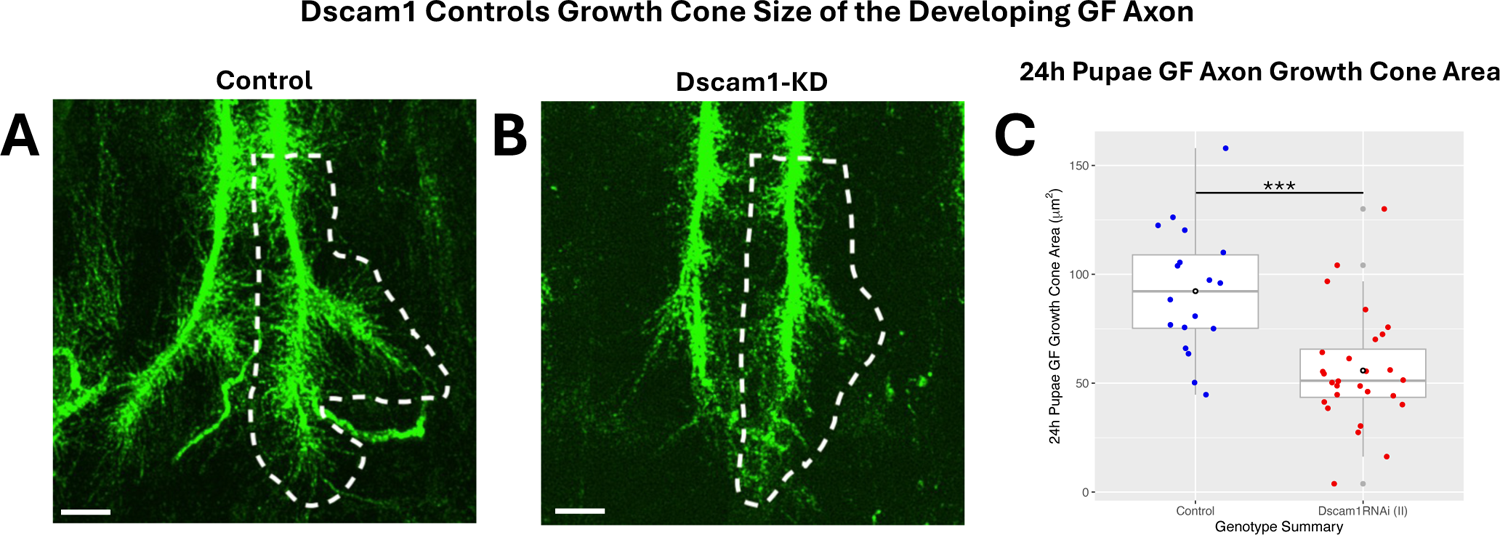
Dscam1 regulates growth cone size in developing GF axons. A-B, Representative maximum intensity projection (MIP) confocal stacks showing GF axon terminals labeled with mCD8::GFP in control (A) and GF-specific Dscam1 knockdown (B) animals using Dscam1RNAi (II), imaged 24 hours after pupation. Filopodial processes are visible at the distal axon tips. The region of interest (ROI) used for growth cone area quantification is outlined in white. Scale bar = 5 μm. C, Quantification of thresholded GFP fluorescence within the ROI shows a significant reduction in growth cone area in Dscam1-KD animals compared to controls (Control: mean = 92.26 μm²; *n* = 18; Dscam1-KD: mean = 55.82 μm²; *n* = 28; *p* = 1.29 × 10⁻⁴; Welch two-sample t-test). Each dot represents a single growth cone; black circle: group mean; grey dots: statistical outliers.

Quantification of growth cone area revealed a significant reduction in Dscam1 knockdown (Dscam1-KD) animals. Dscam1-KD growth cones averaged 55.82 µm², significantly smaller than control growth cones (92.26 µm²; *p* = 1.29 × 10⁻⁴; Fig. 6C). These data indicate that Dscam1 promotes growth cone elaboration during synaptogenesis and that growth cone elaboration is required for subsequent target contact and terminal development.

Together with the structural and functional deficits observed in the mature GF-TTMn circuit, these findings support a model in which Dscam1 is required early in development to ensure adequate growth cone size and downstream synaptic integration.

## Discussion

### Dscam1 regulates synaptic terminal size to support proper circuit function

This study establishes a direct link between presynaptic terminal size and synaptic output in a central circuit and identifies Dscam1 as a critical developmental regulator of this structure-function relationship. In the *Drosophila* escape circuit, knockdown of Dscam1 in the GFs results in longer response latency and a severe reduction in success under repeated high-frequency stimulations, indicating impaired circuit function. Accompanying these deficits are pronounced reductions in the volume of the GF synaptic terminal, the GF-TTMn synapse interface, and the abundance of both electrical (Shak-B) and chemical (Brp) synaptic components. However, the proportion of synaptic proteins relative to terminal or interface size remains constant, implying that Dscam1 loss impairs terminal growth rather than disrupting synaptic protein recruitment or localization. Strong correlations between each anatomical variable alone or measured together, and circuit performance further underscore the importance of presynaptic structure in determining synaptic efficacy. Importantly, we demonstrate that these structural phenotypes emerge during development: at 24 hours after pupation, before mature synapse formation (Lee and Godenschwege, 2015), Dscam1-KD animals already exhibit GF growth defects. Together, these findings suggest that Dscam1 promotes growth cone elaboration and synaptic terminal expansion to support the formation of a properly sized and functional mixed electrochemical synapse.

### Presynaptic terminal size as a determinant of synaptic capacity

Our data support a scaling model in which Dscam1 regulates the size of presynaptic terminals to accommodate appropriate levels of synaptic machinery, including Shak-B gap junctions and Brp-positive active zones. The conserved density and localization of synaptic proteins across genotypes indicate that Dscam1 does not directly regulate their synthesis or placement but instead controls the available presynaptic space, which in turn controls the total amount of protein incorporated. A parallel example has been seen in the regulation of transmitter release at the Drosophila neuromuscular junction (Goel et al., 2019). They showed that one PKC mutant they tested fit a model where transmitter release at the Drosophila neuromuscular junction scaled with the size of the terminal. They too used Brp as their anatomical measure of terminal function and synaptic release as the functional measure and showed that size (in their case, bouton count) scaled with no change in Brp density.

These results highlight a role for Dscam1 in proportional scaling rather than direct regulation of protein localization or transcription. One counter example to proportional scaling is direct control of synaptic proteins at the GF synapse, independent of terminal size. It has been recently demonstrated that the Frazzled gene regulates Shak-B levels at the GF terminal independent of terminal size (Lopez et al., 2025). In this case the intracellular domain of Frazzled appears to regulate the transcription of ShakB directly and in this case ShakB does not scale with size of the terminal. Similarly, regarding direct influence, gap junction recruitment in vertebrate circuits has been proposed to depend on adherens junctions (Cárdenas-García et al., 2024). While homeostatic synaptic scaling has been described in the context of activity-dependent adjustments to synaptic strength (Venkatesan et al., 2020), our findings suggest a distinct form of volumetric scaling, where presynaptic terminal size or metabolic limitation constrains protein abundance. To our knowledge, this represents a novel mechanism of structural constraint at the synapse. Importantly, this mechanism contrasts with models in which trafficking defects or mislocalization of synaptic proteins account for loss-of-function phenotypes (Shapira et al., 2003).

Finally, we find that these presynaptic structural deficits cannot be compensated for by postsynaptic adjustments, which in flies normally contribute to maintaining stable synaptic function (Tripodi et al., 2008).

Dscam1 likely acts early in development to promote growth cone expansion, as we observe reduced growth cone size in Dscam1-KD flies 24 hours after pupation, prior to GF-TTMn mature synapse formation. This suggests that initial morphogenetic events under the control of Dscam1 establish a foundation for subsequent terminal growth and synapse assembly. Whether this effect is mediated through cytoskeletal regulation, modulation of membrane trafficking, or downstream signaling from Dscam1’s intracellular domain remains not yet fully elucidated. Future studies dissecting the molecular pathways downstream of Dscam1 could provide insights into the developmental mechanism that links axon growth to synapse formation.

### Comparison to previous studies of Dscam1 function

Dscam1 has been extensively studied for its roles in axon guidance, dendritic self-avoidance, and branch spacing, particularly in peripheral sensory neurons and developing dendritic arbors (Hughes et al., 2007; Soba et al., 2007; Kim et al., 2013). Our work extends these findings to a well-defined central interneuron with a known physiological output and a precisely mapped synaptic target. We identify a novel role for Dscam1 in controlling the volume of a presynaptic terminal in a central motor circuit, and demonstrate that this structural regulation directly impacts synaptic function and circuit performance.

A prior study concluded that Dscam1 is not required for normal development of the GF axon or dendrites (Wilhelm et al., 2022), based on analyses using the same two RNAi constructs employed here. However, that conclusion was drawn from an assessment limited to gross GF morphology, without examination of the presynaptic terminal. Our findings directly contradict this interpretation: although large-scale neurite patterning appears unaffected, we demonstrate that Dscam1 is in fact essential for fine-scale presynaptic terminal development and circuit-level performance. Thus, the absence of obvious morphological defects at the level of axons or dendrites does not indicate normal synaptic development or function.

In contrast to previous work focusing on neurite patterning or contact-dependent repulsion, our findings suggest that Dscam1 is also essential for building presynaptic compartments large enough to house a full complement of synaptic components. These results are complementary to prior reports describing circuit dysfunction in Dscam1 mutants (Cvetkovska et al., 2013) and suggest a mechanistic basis for such observations rooted in terminal size and synaptic protein constraints. Our dual-mode approach, integrating structure and function in the same animals, further strengthens this interpretation.

### Broader implications and relevance

These findings have broader implications in understanding how neuronal morphology governs circuit connectivity and function. Dscam1 appears to act as a developmental gatekeeper of presynaptic capacity: by enabling sufficient growth cone elaboration and synaptic terminal expansion, it sets the physical limits of synaptic proteins. This scaling mechanism may represent a general principle in neural development, with parallels in other systems where terminal size correlates with synaptic strength or reliability.

More broadly, our data highlight how disruptions in early morphogenetic regulators like Dscam1 can propagate through development to yield dysfunction at the circuit level. Given that human DSCAM is linked to neurodevelopmental disorders, including autism spectrum disorders, bipolar disorder, and Down syndrome (Amano et al., 2008; Narita et al., 2020), and may be genetically linked to late-onset Alzheimer’s (Viard et al., 2022). It is therefore tempting to speculate that similar mechanisms we report here may operate in vertebrate systems. Our findings also raise questions about how neurons compensate for a reduced terminal size of their synaptic partner or altered connectivity and the physiological consequences. Preliminary observations of thinning in the postsynaptic TTMn dendrite following Dscam1-KD suggest potential structural or functional compensation, which merits further investigation.

## Limitations

Despite the robust structure–function relationships observed, this study has several limitations. First, confocal microscopy lacks the resolution to capture ultrastructural details of the synapse interface. Although our volumetric reconstructions are supported by prior correlative EM studies (Hohensee et al., 2008), higher-resolution techniques could more precisely define the spatial relationship between synaptic proteins and membranes. Second, our reliance on RNAi-mediated knockdown may introduce variability in knockdown efficiency. While we validated our phenotypes using two independent RNAi lines, future work could benefit from the quantification of Dscam1 levels following knockdown via mRNA probes or antibodies. Third, our experiments were conducted exclusively in female flies to control for known sex-specific differences in nervous system size and architecture. However, we cannot exclude the possibility of Dscam1-dependent sexual dimorphisms, which warrant future investigation. Lastly, while our data clearly demonstrate that Dscam1 is necessary for proper synapse formation, rescue experiments may help to determine whether restoring terminal size is sufficient to restore function.

### Future directions

Several lines of inquiry emerge from the data we present. First, identifying the intracellular effectors downstream of Dscam1 that control growth cone and synaptic terminal size will be critical. Candidates include regulators of the cytoskeleton and downstream signaling molecules (Worby et al., 2001; Koch et al., 2017; Hernández et al., 2023). While Diap1 overexpression can rescue apoptosis-related defects in some neurodevelopmental models (Singh et al., 2020), it is not known to rescue structural or growth defects resulting from Dscam1 knockdown. Additionally, the similarity between Dscam1 and other attractive guidance receptors, such as Frazzled, invites comparative studies.

Frazzled loss-of-function also produces synaptic defects in the GF circuit, but the phenotypes differ, suggesting distinct but complementary roles in axon guidance and synapse formation (Lopez et al., 2025). Parallel investigation of these guidance receptors could clarify how different guidance cues contribute to final synaptic architecture.

Future studies should also address whether Dscam1 knockdown in the GF alters the morphology or physiology of the postsynaptic TTMn. If dendritic thinning or increased excitability is observed, this could represent a compensatory mechanism. Furthermore, dye-coupling experiments combined with confocal imaging could determine whether changes in gap junction conductance underlie functional impairment. Finally, computational modeling of the GF-TTMn circuit, incorporating precise measures of terminal size and synaptic protein content, could provide predictive insights into how structural perturbations translate to physiological deficit.

In summary, this study identifies Dscam1 as a critical regulator of presynaptic terminal size of the GFs in the *Drosophila* giant fiber system and provides evidence that terminal volume governs synaptic composition and circuit output. These findings offer new perspectives on how early developmental events shape mature nervous system circuit architecture and function.

## Notes

### Competing Interest Statement

The authors have declared no competing interest.

## References

Alavi M, Song M, King GLA, Gillis T, Propst R, Lamanuzzi M, Bousum A, Miller A, Allen R, Kidd T (2016) Dscam1 Forms a Complex with Robo1 and the N-Terminal Fragment of Slit to Promote the Growth of Longitudinal Axons. PLoS Biol 14:e1002560.

Allen Marcus J, Godenschwege TA (2010) Electrophysiological recordings from the Drosophila giant fiber system (GFS). Cold Spring Harb Protoc 2010:pdb.prot5453.

Allen MJ, Godenschwege TA, Tanouye MA, Phelan P (2006) Making an escape: Development and function of the Drosophila giant fibre system. Semin Cell Dev Biol 17:31–41.

Allen Marcus J., Murphey RK (2007) The chemical component of the mixed GF-TTMn synapse in *Drosophila melanogaster* uses acetylcholine as its neurotransmitter. European Journal of Neuroscience 26:439–445.

Allen MJ, Shan X, Caruccio P, Froggett SJ, Moat KG, Murphey RK (1999) Targeted Expression of Truncated *Glued* Disrupts Giant Fiber Synapse Formation in *Drosophila*. The Journal of Neuroscience 19:9374–9384.

Amano K, Yamada K, Iwayama Y, Detera-Wadleigh SD, Hattori E, Toyota T, Tokunaga K, Yoshikawa T, Yamakawa K (2008) Association study between the Down syndrome cell adhesion molecule (DSCAM) gene and bipolar disorder. Psychiatr Genet 18:1–10.

Ammer G, Vieira RM, Fendl S, Borst A (2022) Anatomical distribution and functional roles of electrical synapses in Drosophila. Current Biology 32:2022–2036.e4.

Andrews GL, Tanglao S, Farmer WT, Morin S, Brotman S, Berberoglu MA, Price H, Fernandez GC, Mastick GS, Charron F, Kidd T (2008) Dscam guides embryonic axons by Netrin-dependent and -independent functions. Development 135:3839–3848.

Boerner J, Godenschwege TA (2011) Whole mount preparation of the adult Drosophila ventral nerve cord for giant fiber dye injection. Journal of Visualized Experiments 2–5.

Borgen M, Rowland K, Boerner J, Lloyd B, Khan A, Murphey R (2017) Axon Termination, Pruning, and Synaptogenesis in the Giant Fiber System of *Drosophila melanogaster* Is Promoted by Highwire. Genetics 205:1229–1245.

Cárdenas-García SP, Ijaz S, Pereda AE (2024) The components of an electrical synapse as revealed by expansion microscopy of a single synaptic contact. Elife 13:1–22.

Chen P, Liu Z, Zhang Q, Lin D, Song L, Liu J, Jiao H-F, Lai X, Zou S, Wang S, Zhou T, Li B-M, Zhu L, Pan B-X, Fei E (2022) DSCAM Deficiency Leads to Premature Spine Maturation and Autism-like Behaviors. J Neurosci 42:532–551.

Cvetkovska V, Hibbert AD, Emran F, Chen BE (2013) Overexpression of Down syndrome cell adhesion molecule impairs precise synaptic targeting. Nat Neurosci 16:677–682.

Dascenco D, Erfurth ML, Izadifar A, Song M, Sachse S, Bortnick R, Urwyler O, Petrovic M, Ayaz D, He H, Kise Y, Thomas F, Kidd T, Schmucker D (2015) Slit and Receptor Tyrosine Phosphatase 69D Confer Spatial Specificity to Axon Branching via Dscam1. Cell 162:1140–1154.

Dewa K et al. (2024) Neuronal DSCAM regulates the peri-synaptic localization of GLAST in Bergmann glia for functional synapse formation. Nat Commun 15:458.

Fuerst PG, Bruce F, Tian M, Wei W, Elstrott J, Feller MB, Erskine L, Singer JH, Burgess RW (2009) DSCAM and DSCAML1 Function in Self-Avoidance in Multiple Cell Types in the Developing Mouse Retina. Neuron 64:484–497.

Goel P, Khan M, Howard S, Kim G, Kiragasi B, Kikuma K, Dickman D (2019) A screen for synaptic growth mutants reveals mechanisms that stabilize synaptic strength. Journal of Neuroscience 39:4051–4065.

Hattori D, Chen Y, Matthews BJ, Salwinski L, Sabatti C, Grueber WB, Zipursky SL (2009) Robust discrimination between self and non-self neurites requires thousands of Dscam1 isoforms. Nature 461:644–648.

Hattori D, Millard SS, Wojtowicz WM, Zipursky SL (2008) Dscam-mediated cell recognition regulates neural circuit formation. Annu Rev Cell Dev Biol 24:597–620.

Hernández K, Godoy L, Newquist G, Kellermeyer R, Alavi M, Mathew D, Kidd T (2023) Dscam1 overexpression impairs the function of the gut nervous system in Drosophila. Dev Dyn 252:156–171.

Hohensee S, Bleiss W, Duch C (2008) Correlative electron and confocal microscopy assessment of synapse localization in the central nervous system of an insect. J Neurosci Methods 168:64–70.

Holderith N, Lorincz A, Katona G, Rózsa B, Kulik A, Watanabe M, Nusser Z (2016) Release probability of hippocampal glutamatergic terminals scales with the size of the active zone. Nat Neurosci 19:172–172.

Huang J, Wang Y, Raghavan S, Feng S, Kiesewetter K, Wang J (2011) Human down syndrome cell adhesion molecules (DSCAMs) are functionally conserved with Drosophila Dscam[TM1] isoforms in controlling neurodevelopment. Insect Biochem Mol Biol 41:778–787.

Hughes ME, Bortnick R, Tsubouchi A, Bäumer P, Kondo M, Uemura T, Schmucker D (2007) Homophilic Dscam Interactions Control Complex Dendrite Morphogenesis. Neuron 54:417–427.

Kim JH, Wang X, Coolon R, Ye B (2013) Dscam expression levels determine presynaptic arbor sizes in Drosophila sensory neurons. Neuron 78:827–838.

Koch M, Nicolas M, Zschaetzsch M, de Geest N, Claeys A, Yan J, Morgan MJ, Erfurth M-L, Holt M, Schmucker D, Hassan BA (2017) A Fat-Facets-Dscam1-JNK Pathway Enhances Axonal Growth in Development and after Injury. Front Cell Neurosci 11:416.

Lee LH, Godenschwege TA (2015) Structure-function analyses of tyrosine phosphatase PTP69D in giant fiber synapse formation of Drosophila. Mol Cell Neurosci 64:24–31.

Lopez J, Boerner J, Robbins K, Pena RFO, Murphey R (2025) Frazzled/DCC regulates gap junction formation at a Drosophila giant synapse through transcription. [In press - eNeuro]

Lu Zhongmin, Chouhan AK, Borycz JA, Lu Zhiyuan, Rossano AJ, Brain KL, Zhou Y, Meinertzhagen IA, Macleod GT (2016) High-Probability Neurotransmitter Release Sites Represent an Energy-Efficient Design. Current Biology 26:2562–2571.

Matthews BJ, Grueber WB (2011) Dscam1-mediated self-avoidance counters netrin-dependent targeting of dendrites in Drosophila. Curr Biol 21:1480–1487.

Matthews BJ, Kim ME, Flanagan JJ, Hattori D, Clemens JC, Zipursky SL, Grueber WB (2007) Dendrite Self-Avoidance Is Controlled by Dscam. Cell 129:593–604.

Millard SS, Lu Z, Zipursky SL, Meinertzhagen IA (2010) Drosophila Dscam Proteins Regulate Postsynaptic Specificity at Multiple-Contact Synapses. Neuron 67:761–768.

Montesinos ML (2014) Roles for DSCAM and DSCAML1 in central nervous system development and disease. Adv Neurobiol 8:249–270.

Narita A et al. (2020) Clustering by phenotype and genome-wide association study in autism. Transl Psychiatry 10:290.

Phelan P, Nakagawa M, Wilkin M, Moffat K, O’Kane C, Davies J, Bacon J (1996) Mutations in shaking-B prevent electrical synapse formation in the Drosophila giant fiber system. The Journal of Neuroscience 16:1101–1113.

Rollenhagen A, Lübke JHR (2006) The morphology of excitatory central synapses: from structure to function. Cell Tissue Res 326:221–37.

Sachse SM et al. (2019) Nuclear import of the DSCAM -cytoplasmic domain drives signaling capable of inhibiting synapse formation. EMBO J 38.

Schmucker D, Clemens JC, Shu H, Worby CA, Xiao J, Muda M, Dixon JE, Zipursky SL (2000) Drosophila Dscam Is an Axon Guidance Receptor Exhibiting Extraordinary Molecular Diversity. Cell 101:671–684.

Shapira M, Zhai RG, Dresbach T, Bresler T, Torres VI, Gundelfinger ED, Ziv NE, Garner CC (2003) Unitary Assembly of Presynaptic Active Zones from Piccolo-Bassoon Transport Vesicles. Neuron 38:237–252.

Singh MD, Jensen M, Lasser M, Huber E, Yusuff T, Pizzo L, Lifschutz B, Desai I, Kubina A, Yennawar S, Kim S, Iyer J, Rincon-Limas DE, Lowery LA, Girirajan S (2020) NCBP2 modulates neurodevelopmental defects of the 3q29 deletion in Drosophila and Xenopus laevis models. PLoS Genet 16:e1008590.

Soba P, Zhu S, Emoto K, Younger S, Yang S-J, Yu H-H, Lee T, Jan LY, Jan Y-N (2007) Drosophila Sensory Neurons Require Dscam for Dendritic Self-Avoidance and Proper Dendritic Field Organization. Neuron 54:403–416.

Sun W, You X, Gogol-Döring A, He H, Kise Y, Sohn M, Chen T, Klebes A, Schmucker D, Chen W (2013) Ultra-deep profiling of alternatively spliced Drosophila Dscam isoforms by circularization-assisted multi-segment sequencing. EMBO J 32:2029–2038.

Tripodi M, Evers JF, Mauss A, Bate M, Landgraf M (2008) Structural homeostasis: Compensatory adjustments of dendritic arbor geometry in response to variations of synaptic input. PLoS Biol 6:2172–2187.

Uthaman SB, Godenschwege TA, Murphey RK (2008) A mechanism distinct from highwire for the Drosophila ubiquitin conjugase bendless in synaptic growth and maturation. J Neurosci 28:8615–8623.

Van Vactor D, Sigrist SJ (2017) Presynaptic morphogenesis, active zone organization and structural plasticity in Drosophila. Curr Opin Neurobiol 43:119–129.

Venkatesan S, Subramaniam S, Rajeev P, Chopra Y, Jose M, Nair D (2020) Differential Scaling of Synaptic Molecules within Functional Zones of an Excitatory Synapse during Homeostatic Plasticity. eNeuro 7:ENEURO.0407-19.2020.

Viard J et al. (2022) Chr21 protein-protein interactions: enrichment in proteins involved in intellectual disability, autism, and late-onset Alzheimer’s disease. Life Sci Alliance 5.

Wilhelm N, Kumari S, Krick N, Rickert C, Duch C (2022) Dscam1 Has Diverse Neuron Type-Specific Functions in the Developing Drosophila CNS. eNeuro 9.

Williams DL, Sikora VM, Hammer MA, Amin S, Brinjikji T, Brumley EK, Burrows CJ, Carrillo PM, Cromer K, Edwards SJ, Emri O, Fergle D, Jenkins MJ, Kaushik K, Maydan DD, Woodard W, Clowney EJ (2022) May the Odds Be Ever in Your Favor: Non-deterministic Mechanisms Diversifying Cell Surface Molecule Expression. Front Cell Dev Biol 9.

Worby CA, Simonson-Le N, Clemens JC, Kruger RP, Muda M, Dixon JE (2001) The Sorting Nexin, DSH3PX1, Connects the Axonal Guidance Receptor, Dscam, to the Actin Cytoskeleton. Journal of Biological Chemistry 276:41782–41789.

